# Epstein–Barr Virus BALF0/1 Subverts the Caveolin and ERAD Pathways to Target B-cell Receptor Complexes for Degradation

**DOI:** 10.1101/2024.01.04.574276

**Authors:** Stephanie Pei Tung Yiu, Michael Weekes, Benjamin E. Gewurz

## Abstract

Epstein–Barr virus (EBV) establishes persistent infection, causes infectious mononucleosis, is a major trigger for multiple sclerosis and contributes to multiple cancers. Yet, knowledge remains incomplete about how the virus remodels host B cells to support lytic replication. We previously identified that EBV lytic replication results in selective depletion of plasma membrane B-cell receptor (BCR) complexes, comprised of immunoglobulin and the CD79A and CD79B signaling chains. Here, we used proteomic and biochemical approaches to identify that the EBV early lytic protein BALF0/1 is responsible for EBV lytic cycle BCR degradation. Mechanistically, an immunoglobulin heavy chain cytoplasmic tail KVK motif was required for ubiquitin-mediated BCR degradation, while CD79A and CD79B were dispensable. BALF0/1 subverted caveolin-mediated endocytosis to internalize plasma membrane BCR complexes and to deliver them to the endoplasmic reticulum. BALF0/1 stimulated immunoglobulin heavy chain cytoplasmic tail ubiquitination, which together with the ATPase valosin-containing protein/p97 drove ER-associated degradation of BCR complexes by cytoplasmic proteasomes. BALF0/1 knockout reduced the viral load of secreted EBV particles from B-cells that expressed a monoclonal antibody against EBV glycoprotein 350 and increased viral particle immunoglobulin incorporation. Consistent with downmodulation of plasma membrane BCR, BALF0/1 overexpression reduced viability of a diffuse large B-cell lymphoma cell line dependent upon BCR signaling. Collectively, our results suggest that EBV BALF0/1 downmodulates immunoglobulin upon lytic reactivation to block BCR signaling and support virion release.

**SIGNIFICANCE:** EBV uses a biphasic lifecycle, in which it switches between a latent state that facilitates immune evasion and a lytic state, where virion are secreted. However, when EBV infects a B-cell that makes antibody against a virion protein, EBV must have a strategy to escape becoming trapped, since maturing virion and antibody each traffic through the secretory pathway. We identified that an EBV-encoded protein expressed, BALF0/1, associates with and targets immunoglobulin complexes for degradation. Intriguingly, BALF0/1 subverts the caveolin-1 and ERAD pathways to route antibody from the plasma membrane to cytoplasmic proteasomes for degradation. We present evidence that this enhances EBV secretion from cells that produce antibody against a viral glycoprotein, which could otherwise trap virus.

## INTRODUCTION

Epstein–Barr virus (EBV) establishes lifelong infection in >95% of adults worldwide, is the major viral trigger for multiple sclerosis and is associated with ∼2% of human cancers (1, 2). These include endemic Burkitt lymphoma, Hodgkin lymphoma, post-transplant and HIV-associated lymphomas, T and NK-cell lymphomas, nasopharyngeal and gastric carcinomas (3, 4). EBV uses a biphasic lifecycle to colonize the B-cell compartment, to spread between B and epithelial cells, and to transmit between hosts (5–7). The EBV lytic cycle is triggered by the immediate early genes BZLF1 and BRLF1, which induce expression of 35 early genes important for viral DNA replication and then 40 late genes that include structural proteins important for production of infectious virion. Much remains to be learned about how EBV subverts host innate and adaptive immune barriers in order to establish latency, reactivate within the heart of the adaptive immune system.

EBV has evolved multiple mechanisms to interfere with antigen presentation pathways, thereby suppressing cell-mediated immune responses (8–15). However, elevated antibody titers against viral antigens are found in numerous EBV-associated disease states, including with infectious mononucleosis, endemic Burkitt lymphoma, nasopharyngeal carcinoma, classical Hodgkin Lymphoma, and multiple sclerosis (16–18). While Epstein-Barr virion evasion of complement activation has been described (PMID 2844953), viral strategies to protect virion from immunoglobulin, including that produced within the infected cell where viral particles mature, are less well-studied.

To gain insights into how EBV remodels host B-cells in support of lytic replication, we used tandem-mass-tag-based MS3 mass spectrometry to perform quantitative temporal proteomic analysis in EBV+ Burkitt lymphoma B cells, prior to and at four time points following lytic reactivation (19). Unexpectedly, this analysis revealed that B-cell receptor (BCR) complexes, comprised of immunoglobulin (Ig) heavy chain, light chain and non-covalently associated CD79A/B (also known as Igα/β) signaling chains, were selectively and highly depleted upon EBV lytic reactivation. In IgM expressing cells, the secretory IgM joining-chain (J-chain) was also depleted, whereas only a small number of host cell proteins exhibited significant downmodulation, suggesting selective post-translational targeting (19). Whereas activated BCR complexes are typically delivered to lysosomes to support presentation of peptide antigens to CD4+ T-cells by major histocompatibility complex (MHC) class II molecules (20), EBV instead routes BCR to cytosolic proteasomes by an unidentified pathway (19). Which EBV gene targets BCR complexes for degradation, and how it targets BCR complexes embedded within B-cell membranes to cytosolic proteasomes has remained unknown. However, acyclovir treatment did not prevent EBV lytic cycle BCR depletion, indicating that viral late lytic genes are not required (19). Here, we used a recently constructed EBV lytic cycle protein-protein interaction map (21) to generate the hypothesis that the EBV early lytic cycle encoded BALF0/1 targets BCR complexes for proteasomal degradation. Focused over-expression and CRISPR studies supported and extended the hypothesis, demonstrating that BALF0/1 subverts caveolar endocytosis and ER-associated degradation pathways to target BCR for degradation.

## RESULTS

### BALF0/1 targets the B-cell receptor for degradation during EBV lytic cycle

To gain insights into how EBV targets the BCR for cytoplasmic proteasome degradation, we interrogated a recently generated EBV protein-protein interaction network map, generated in Burkitt lymphoma cells induced for EBV lytic replication (21). Our proteomic analysis identified high-confidence interactions between the immunoglobulin (Ig) heavy chain and two EBV early lytic proteins, BALF0/1 and BARF1 (Figure 1A). BALF0/1 associated with multiple proteasome components, with multiple proteins involved in microtubule associated protein transport, and with B-cell receptor-associated protein 31 (BCAP31), a chaperone that plays a role in the ER associated degradation (ERAD) pathway (Figure 1A). Of note, two in-frame methionine codons can be used to generate a 220 residue BALF0 or a 189 residue BALF1 protein (22). BALF0/1 is predicted to exhibit structural homology with MCL1 (<u>Figure S1</u>A). Similarly, proteomic analysis highlighted interaction between BARF1 and several ubiquitin-proteasome pathway proteins (Figure 1A)(21). BARF1 exhibits homology with the colony stimulating factor receptor and also has limited amino acid sequence similarity with CD80 (11, 23).

**Figure 1.**
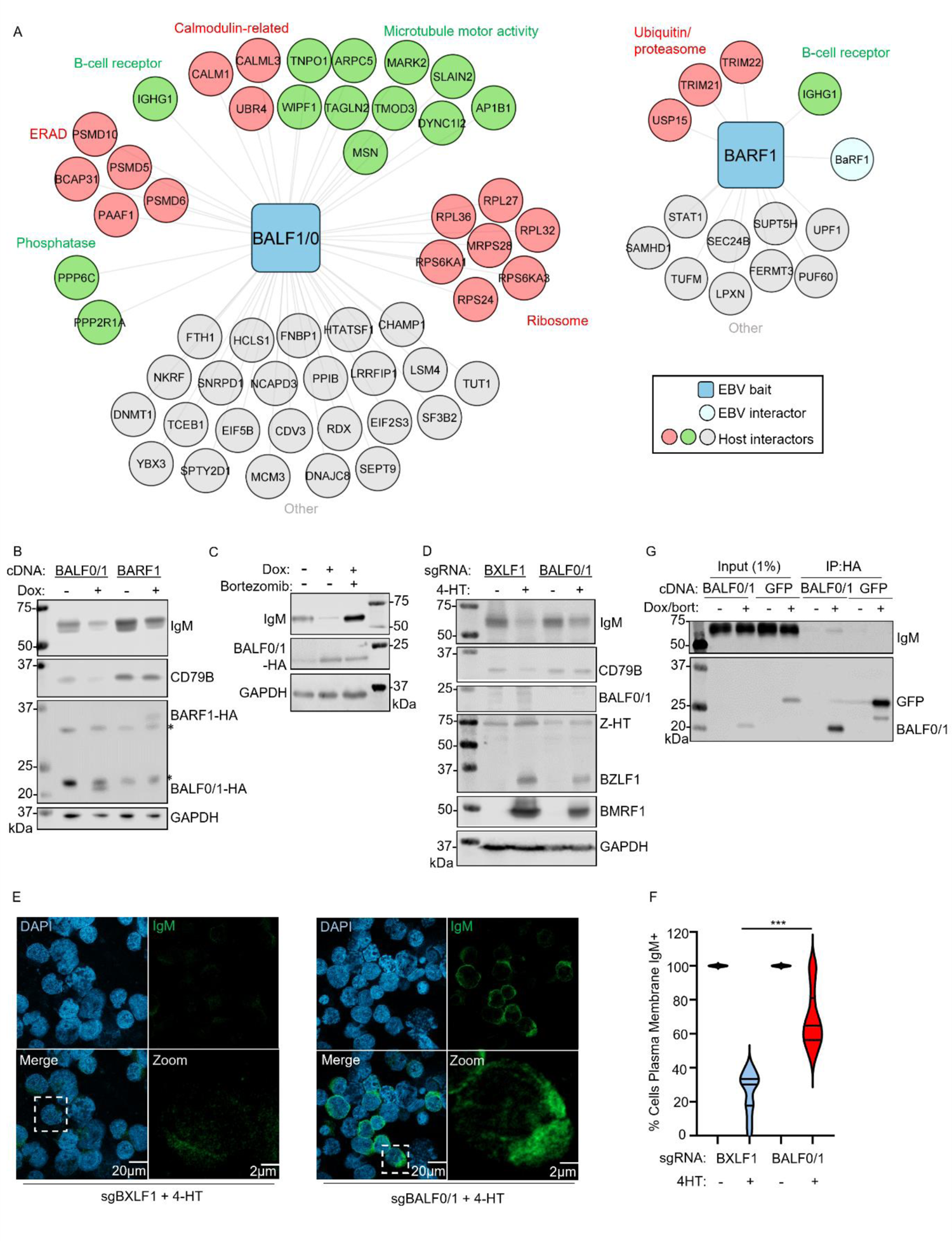
EBV early lytic protein BALF0/1 targets the B-cell receptor complex for degradation (A) EBV B-cell lytic cycle BALF0/1 (left) and BARF1 (right) protein-protein interaction network, adapted from (21). DAVID was used to identify enriched pathways among BALF0/1 interactors. (B) Immunoblot analysis of whole cell lysates (WCL) from latent P3HR-1 Burkitt cells induced for expression of BALF0/1 or BARF1 cDNAs by 5 µg/ml doxycycline (Dox) for 24 h, as indicated. * indicates background band. (C) Immunoblot analysis of WCL from P3HR-1 cells expressing BALF0/1 cDNA, induced by 5 µg/ml doxycycline (Dox) in the presence or absence of 5 nM of the proteasome inhibitor bortezomib for 24 h, as indicated. (D) Immunoblot analysis of WCL from Cas9+ P3HR-1 ZHT/RHT cells expressing the indicated sgRNA, induced into lytic replication by 400 nM 4-HT for 24 h. (E) Immunofluorescence analysis of IgM in Cas9+ P3HR-1 ZHT/RHT cells expressing the indicated BXLF1 or BALF0/1 targeting sgRNA. Cells were 4-HT induced for lytic replication for 24 h. See also Fig. S1B. (F) Mean ± standard error of the mean (SEM) percentage of cells with detectable IgM signal as in panel E and <u>Figure S1</u>B of P3HR-1 cells expressing the indicated BXLF1 or BALF0/1 sgRNA, 4-HT induced as indicated, using data from 15 randomly selected panels of 300 cells from n=3 replicates, analyzed using the ImageJ ComDet plugin. (G) Immunoblot analysis of 1% input and anti-HA immunopurified from P3HR-1 stably expressing BALF0/1-HA or GFP-HA, treated with 5 µg/ml doxycycline and 5 nM bortezomib (bort) for 24 h, as indicated. Statistical analysis was performed with Student’s t-test unless otherwise specified. ***p < 0.001. White bars indicate scale. See also <u>Figure S</u>1. Blots are representative of at least n=2 replicates

To test whether either BALF0/1 or BARF1 can independently deplete BCR components, we conditionally expressed cDNA encoding either of these two EBV early genes in P3HR-1 Burkitt lymphoma cells. Expression of BALF0/1, but not BARF1, depleted both CD79B and IgM heavy chains, suggesting that BALF0/1 can trigger BCR depletion even in the absence of an EBV lytic cycle (Figure 1B). Since we previously found that the EBV lytic cycle routes the BCR to cytosolic proteasomes, we next measured BCR subunit abundances in BALF0/1+ P3HR-1 cells, in the absence or presence of the proteasome inhibitor bortezomib (19). Bortezomib increased IgM heavy chain abundance in BALF0/1 expressing cells (Figure 1C).

To build on these results, we next asked whether BALF0/1 was necessary for BCR loss in B-cells triggered for lytic reactivation. For this analysis, we used Cas9+ P3HR-1 cells with a conditional EBV lytic reactivation system, where the EBV immediate early BZLF1 and BRLF1 alleles are fused to a mutant estrogen receptor binding domain (24, 25). Addition of 4-hydroxytamoxifen (4-HT) triggers nuclear translocation of the BZLF1 and BRLF1 chimeras, termed ZHT and RHT, respectively, to drive EBV early gene expression. To establish CRISPR-edited cells, single guide RNAs (sgRNAs) were separately expressed to target *BALF0/1* or instead as a control to target the EBV early lytic gene *BXLF1,* which encodes the viral thymidine kinase that is dispensable for EBV replication (26, 27). BALF0/1 or BXLF1 knockout (KO) did not perturb expression of the endogenous immediate early BZLF1 or early BMRF1 genes upon 4-HT addition, indicating fidelity of EBV genomes in CRISPR-edited P3HR-1 cells. Consistent with our over-expression analysis, *BALF0/1,* but not *BXLF1* KO significantly increased IgM and CD79B abundances in lytic P3HR-1 cells (Figure 1D-F and <u>S1</u>B). Furthermore, we validated that IgM heavy chains co-immunoprecipitated with BALF0/1 from whole cell lysates of bortezomib-treated P3HR-1 cells induced for lytic replication (Figure 1G). Collectively, our results suggest that BALF0/1 associates with and destabilizes BCR complexes, beginning in the early phase of EBV lytic replication.

### BALF0/1 utilizes the caveolae pathway to internalize plasma membrane BCR

We next investigated how cytoplasmic BALF0/1 routes membrane embedded BCR complexes, including the large pool of plasma membrane BCR, to cytoplasmic proteasomes. We hypothesized that BALF0/1 must first trigger BCR endocytosis prior to its degradation, given that we previously observed intracellular IgM heavy chain in bortezomib-treated cells triggered for lytic replication (19). To further characterize this process, we performed confocal microscopy on 4-HT and bortezomib treated P3HR-1 cells expressing control BXLF1 or BALF0/1 sgRNAs. Consistent with our prior observations, internalized IgM puncta were evident in bortezomib-treated lytic BXLF1 KO control cells, but remained largely localized at the plasma membrane in BALF0/1 edited cells induced for replication (Figure 2A-B). Moreover, BALF0/1 colocalized with internalized IgM in most bortezomib-treated P3HR-1 cells and in with internalized IgM in rare cells in which IgM was not depleted in the absence of bortezomib treatment (Figure 2C). As BALF0/1 is not known to insert into the plasma membrane, this result suggests that it triggers BCR endocytosis.

**Figure 2.**
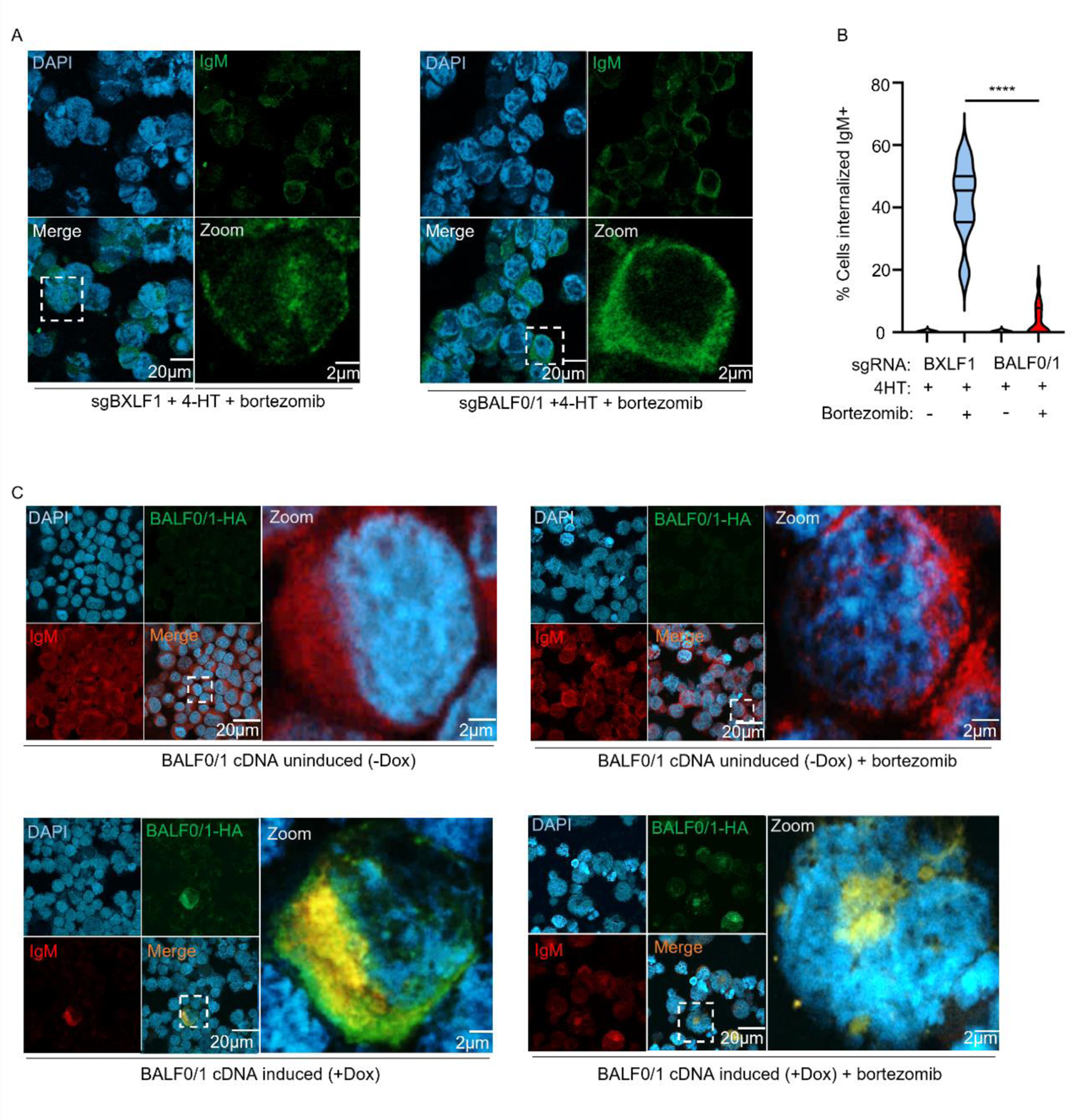
BALF0/1 triggers endocytosis of the BCR complex (A) Immunofluorescence analysis of IgM subcellular distribution in Cas9+ P3HR-1 cells expressing the indicated BXLF1 or BALF0/1 targeting sgRNA. Cells were 4-HT induced for lytic replication and treated with bortezomib for 24 h. Magnification of the white boxed areas are shown in the bottom right of each panel. (B) Mean ± SEM percentage of cells with internalized IgM as in panel A and Figure 1E of P3HR-1 cells expressing the indicated BXLF1 or BALF0/1 sgRNA, 4-HT induced in the presence of bortezomib, as indicated, using data from 15 randomly selected panels of 300 cells from n=3 replicates, analyzed using the ImageJ ComDet plugin. (C) Immunofluorescence analysis of IgM and BALF0/1 colocalization, as determined by staining for IgM (red) and for HA-tagged BALF0/1 (green) in P3HR-1 cells, mock-induced or induced for BALF0/1-HA expression by doxycycline addition in the absence or presence of bortezomib for 24 h. Yellow indicates area of overlap in the merged images. Magnified images of the white boxed areas are shown to the right of each panel. Statistical analysis was performed with Student’s t-test unless otherwise specified. ****p < 0.0001. White bars indicate scale.

Caveolar and clathrin mediated endocytosis are the two major pathways that internalize plasma membrane receptors. Clarthin pathway cargo are predominantly recycled or routed to lysosomes for degradation (28), whereas caveolar pathway cargo travel retrograde to the Golgi, ER or to endosomes (29–33). Since our proteomic analysis also identified high confidence BALF0/1 associations with proteins related to calmodulin and microtubule motor activity (Figure 1A), we next investigated whether either pathway is required for BALF0/1 downmodulation of BCR complexes. P3HR-1 were treated with chlorpromazine or genistein, which inhibit the clarthin or caveolae endocytosis pathways, respectively, together with 4-HT. Interestingly, genistein but not chlorpromazine impaired plasma membrane IgM down-modulation and BCR degradation in a dose-dependent manner upon EBV lytic reactivation (Figure 3A-C and <u>S2</u>A-B). However, since genistein also reduced EBV lytic cycle protein expression at elevated doses (<u>Figure S2</u>B), we further validated this result by expressing BALF0/1 cDNA in latent P3HR-1 cells, in the absence or presence of genistein. Genistein rescued BCR expression upon BALF0/1 expression without downmodulating BALF0/1 levels (Figure 3B). Moreover, CRISPR depletion of the key caveolae structural protein caveolin-1 (CAV-1), which is required for caveolin-mediated internalization, also impaired IgM degradation upon EBV lytic reactivation (Figure 3D). CAV-1 depletion also increased plasma membrane IgM abundance in cells induced for lytic replication (Figure 3D-E and <u>S2</u>C). In control cells induced for lytic replication, bortezomib resulted in the accumulation of intracellular IgM that partially overlapped with the endoplasmic reticulum (ER) resident protein calnexin. By contrast, IgM exhibited plasma membrane distribution and had little co-localization with calnexin in CRISPR CAV-1 depleted cells treated with 4-HT and bortezomib (Figure 3E-F and <u>S2</u>D). These results suggest that BALF0/1 triggers plasma membrane BCR internalization via the caveolin pathway.

**Figure 3.**
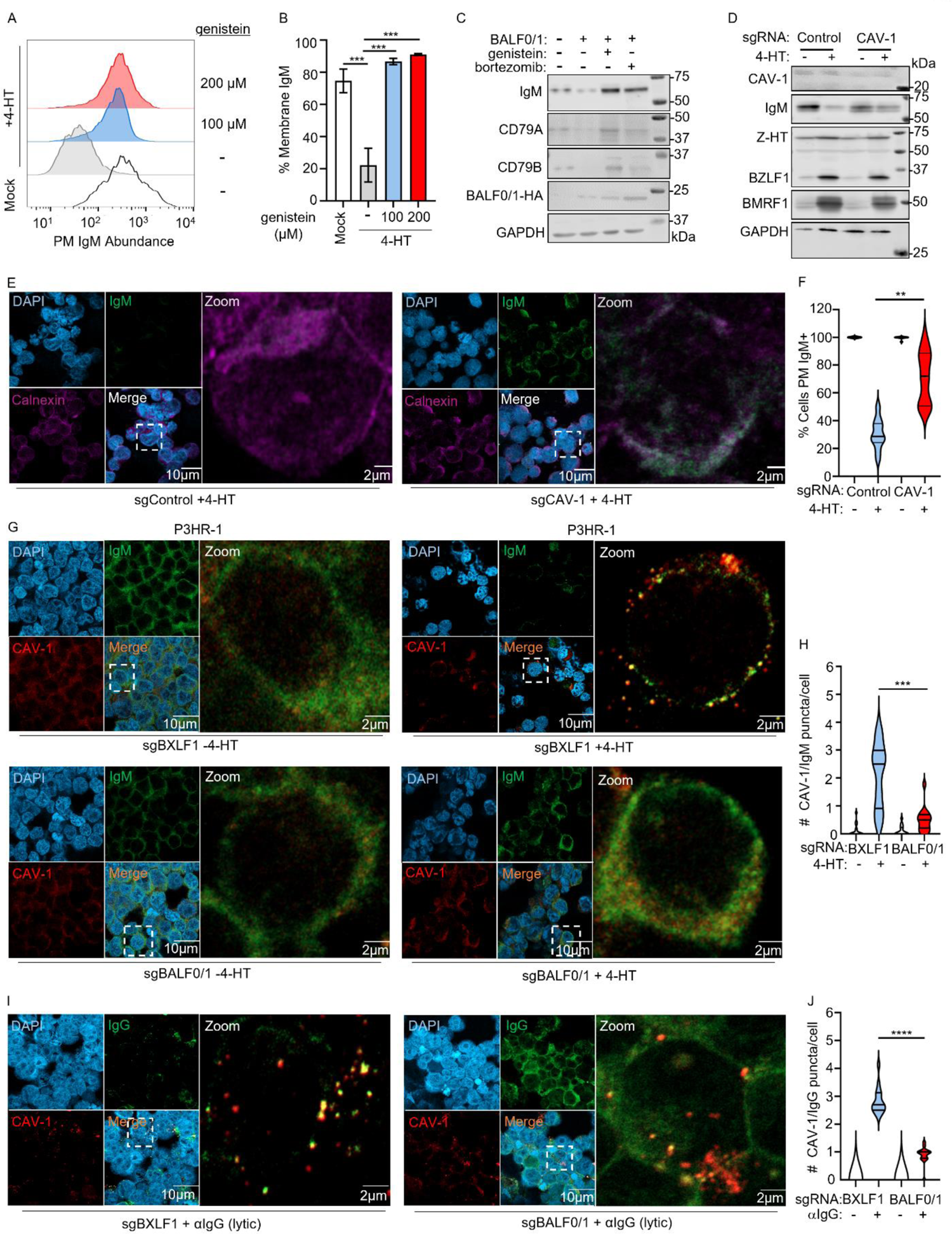
BALF0/1 triggers BCR endocytosis via the caveolae pathway (A) FACS analysis of plasma membrane (PM) IgM levels in P3HR-1 cells induced for lytic replication by 4HT, in the absence or presence of the indicated concentration of genistein for 24h, as indicated. (B) Mean ± SEM of PM IgM abundances from n=3 independent replicates, as in panel A. (C) Immunoblot analysis of WCL from P3HR-1 cells induced for BALF0/1 expression by 5 µg/ml doxycycline in the absence or presence of 5 nM bortezomib or 100 µM genistein for 24 h, as indicated. (D) Immunoblot analysis of WCL from P3HR-1 cells expressing control or caveolin-1 (CAV-1) targeting sgRNA and 4-HT induced into lytic replication. (E) Immunofluorescence analysis of IgM expression in P3HR-1 cells expressing control or CAV-1 targeting sgRNA, induced for lytic replication by 4-HT treatment for 24 h. For comparison, ER-resident calnexin is shown. (F) Mean ± SEM percentage of cells with PM IgM expression as in panel E and <u>Figure S2</u>C of P3HR-1 cells expressing the indicated control or CAV-1 sgRNA, 4-HT induced, as indicated, using data from 12 randomly selected panels of 240 cells from n=3 replicates, analyzed using the ImageJ ComDet plugin. (G) Immunofluorescence analysis of IgG and CAV-1 subcellular localization in Cas9+ P3HR1 ZHT/RHT cells expressing BXLF1 or BALF0/1 sgRNA, as indicated. Cells were induced into lytic cycle by 4HT crosslinking for 16h, as indicated. (H) Mean ± SEM number of cells with co-localized IgM and CAV-1 puncta as in panel G of P3HR-1 cells expressing the indicated BXLF1 or BALF0/1 sgRNA, uninduced or 4-HT induced, as indicated, using data from 20 randomly selected panels of 400 cells from n=3 replicates, analyzed using the ImageJ ComDet plugin. (I) Immunofluorescence analysis of CAV-1 and IgG subcellular distributions in EBV+ Cas9+ Akata expressing BXLF1 or BALF0/1 targeting sgRNA and induced for lytic replication by α-IgG crosslinking for 48 h. (J) Mean ± SEM number of cells with co-localized IgG and CAV-1 signals as in panel I and <u>Figure</u> <u>S3</u> of EBV+ Cas9+ Akata cells expressing the indicated BXLF1 or BALF0/1 sgRNA, α-IgG crosslinked, as indicated, using data from 20 randomly selected panels of 400 cells from n=3 replicates, analyzed using the ImageJ ComDet plugin. Statistical analysis was performed with Student’s t-test unless otherwise specified. ****p < 0.0001. ***p < 0.001. **p < 0.01. White bars indicate scale. See also <u>Figure S</u>2-3. Blots are representative of at least n=2 replicates.

Consistent with a major Cav-1 role downstream of BALF0/1, 4-HT lytic reactivation triggered extensive CAV-1 puncta formation in control BXLF1 KO, but to a lesser extent in BALF0/1 KO P3HR-1 (Figure 3G-H). Interestingly, anti-IgG crosslinking, which triggers EBV lytic reactivation, also induced Cav-1 puncta formation in Akata Burkitt B-cells, both in EBV+ and EBV-subclones (34) (<u>Figure S2</u>E). However, IgG puncta more frequently co-localized with CAV-1 in EBV+ Akata cells (<u>Figure S2</u>E). This result suggests that the EBV lytic cycle, rather than IgG cross-linking alone, induces IgG internalization into CAV-1-containing vesicles at the 48 hour timepoint post-Ig crosslinking. Furthermore, CAV-1 puncta more frequently co-localized with residual IgG signals in Ig-crosslinked BXLF1 edited control Akata cells than in BALF0/1 edited Akata cells, even though they were large depleted relative to levels observed in BALF0/1 depleted cells (Figure 3I-J and <u>S3</u>). Collectively, these data support a model by which BALF1 subverts the caveolin pathway to endocytose plasma membrane BCR complexes.

### BALF0/1 uses the ER-associated degradation pathway to route BCR to proteasomes

We next hypothesized that BILF1 routes internalized BCR complexes to the ER in order use ER-associated degradation (ERAD) (35–37) to reach cytoplasmic proteasomes, in a manner reminiscent of the cholera toxin cell entry pathway (38). In support, our proteomic analysis identified high confidence BALF0/1 interactions with B-cell receptor-associated protein 31 (BCAP31, also called BAP31), an ER resident protein with roles in quality control, including in ERAD (39–43) and with multiple 26S proteosome subunits (Figure 1A). To test this hypothesis, we first performed confocal microscopy on latent vs reactivated P3HR-1, in the absence or presence of bortezomib to stabilize BCR in reactivated cells. Whereas IgM and the ER-resident protein calnexin did not significantly colocalize in latent cells, likely because BCR rapidly transits through the ER on route to the plasma membrane, 4-HT treatment caused near total loss of IgM (Figure 4A, top right panel). Interestingly, in bortezomib-treated cells, 4-HT instead triggered IgM internalization, large regions of which co-localized with the ER-resident chaperone calnexin (Figure 4A-C). Similar calnexin/IgG colocalization was observed in bortezomib treated EBV+ Akata cells triggered for lytic reactivation by IgG-crosslinking (<u>Figure S4</u>A-C). We note that not all 4-HT or anti-IgG treated cells reactivate EBV, and that not all BCR aggregates were captured in a 2-dimensional slice, potentially resulting in an under-estimate of the extent of EBV lytic cycle induced Ig/calnexin colocalization. Our results suggest that BALF0/1 routes internalized BCR to the ER, and that proteasome inhibition causes a BCR population to be retained within the ER.

**Figure 4.**
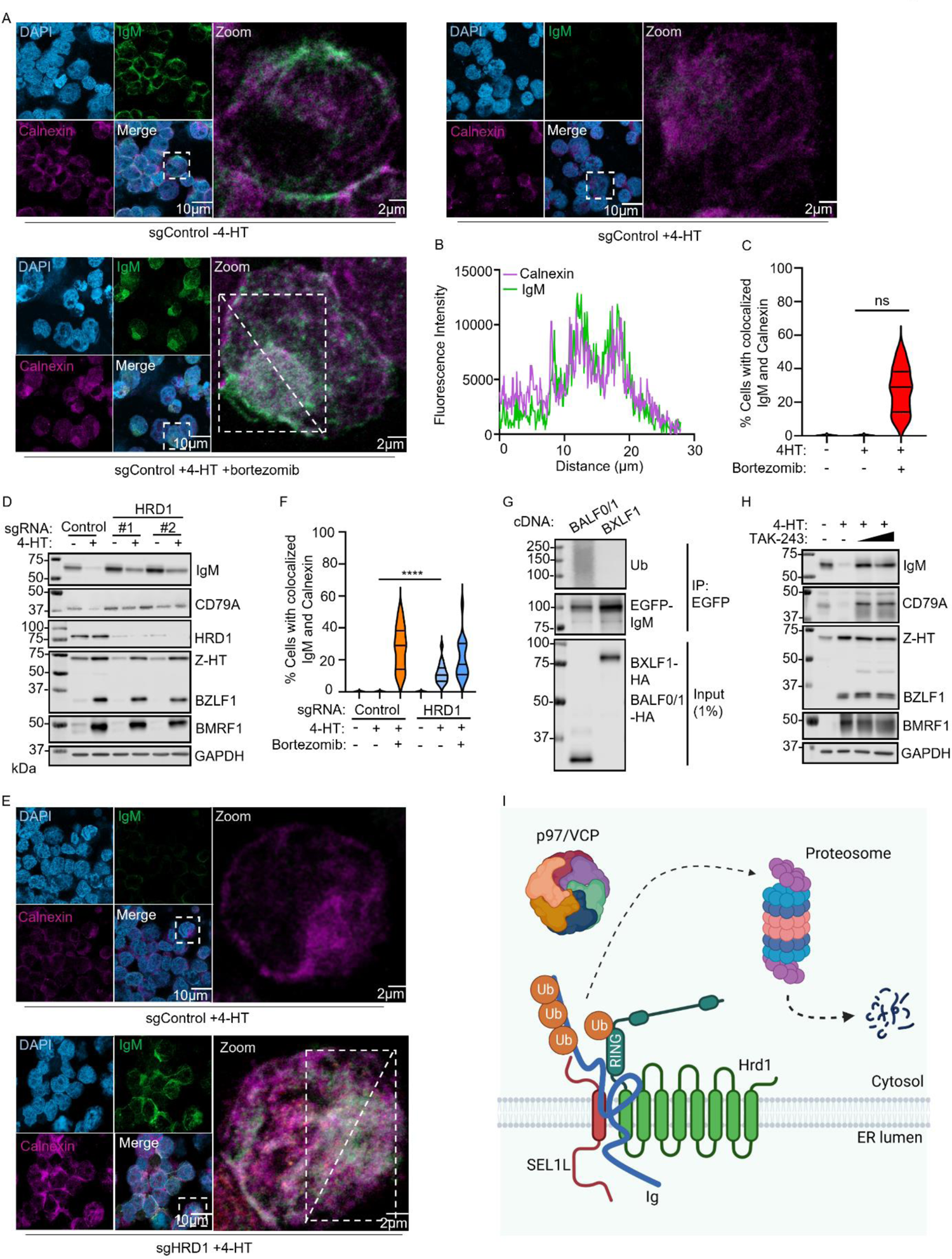
BALF1/0 subverts the ER-associated degradation pathway to route BCR to cytosolic proteasomes (A) Immunofluorescence analysis of IgM and calnexin subcellular distribution in Cas9+ P3HR-1 cells expressing control sgRNA and 4-HT induced for lytic replication for 24 h, in the absence or presence of bortezomib. (B) Line scanning of fluorescence intensity for calnexin (magenta) and IgM (green) in the annotated the white rectangle shown in panel A. (C) Mean ±SEM percentage of cells with overlapping calnexin and IgM signals as in panel A of P3HR-1 cells expressing the indicated control sgRNA, 4-HT induced in the absence or presence of bortezomib as indicated, using data from 12 randomly selected panels of 240 cells from n=3 replicates, analyzed using the ImageJ ComDet plugin. (D) Immunoblot analysis of WCL from Cas9+ P3HR-1 cells expressing the indicated HRD1-targeting sgRNA, 4-HT induced into lytic cycle for 24 h. Representative of n=2 replicates. (E) Immunofluorescence analysis of IgM and calnexin subcellular distribution in Cas9+ P3HR-1 cells expressing control or HRD1 sgRNA, as indicated and 4-HT induced for lytic replication for 24 h. See also Fig. S5D. (F) Mean ±SEM percentage of cells with overlapping calnexin and IgM signals as in panel E and <u>Figure S5</u>B-D of P3HR-1 cells expressing the indicated control or HRD1 sgRNA, uninduced or 4-HT induced in the absence or presence of bortezomib as indicated, using data from 15 randomly selected panels of 300 cells from n=3 replicates, analyzed using the ImageJ ComDet plugin. (G) Immunoblot analysis of 1% input and anti-EGFP immunopurified complexes from 293T cells transiently expressing EGFP-IgM heavy chain chimera together with BALF0/1 or BXLF1 and blotted for poly-ubiquitin chains, for EGFP or for HA, as indicated. (H) Immunoblot analysis of WCL from P3HR-1 cells 4-HT induced into lytic cycle for 24 h, in the absence or presence of increasing doses of the ubiquitin E1 enzyme inhibitor TAK-243 (1 and 2 µM). (I) Schematic model of immunoglobulin (Ig) dislocation via the ERAD pathway, where the Ig heavy chain is ubiquitinated and dislocated from the ER to the cytosol in a HRD1 and p97/VCP dependent manner, where it is degraded by proteasomes. Statistical analysis was performed with Student’s t-test unless otherwise specified. ****p < 0.0001. White bars indicate scale. See also <u>Figure S</u>4-5. Blots are representative of at least n=2 independent experiments.

To further examine ERAD roles in BALF0/1-mediated BCR down-regulation, we depleted Hrd1 (HMG-CoA reductase degradation protein 1), an ER-resident E3 ubiquitin ligase that forms an ER-membrane spanning channel important for ERAD (44–46) or of BALF1/0 associated BCAP31. HRD1 KO diminished the extent of IgM and CD79A loss observed in P3HR-1 induced for lytic reactivation, despite having no appreciable effects on immediate early BZLF1 or early gene BMRF1 expression (Figure 4D). Similarly, the small molecule ERAD inhibitor eeyarestatin I (47), which blocks AAA ATPase p97/VCP mediated cytoplasmic shuttling of ERAD cargo (48), significantly impaired BCR degradation in a dose-dependent manner, albeit with a degree of EBV lytic cycle gene repression. By contrast, the lysosomal degradation inhibitors leupeptin, E64 or bafilomycin failed to rescue EBV lytic cycle BCR degradation (<u>Figure S5</u>A). Furthermore, we observed calnexin-colocalized BCR aggregates in a significant number of P3HR-1 Hrd1 KO cells induced for lytic replication, which accumulated to a similar degree as in bortezomib-treated cells (Figure 4E-F and <u>S5</u>B-D). Despite its association with BALF0/1, BCAP31 KO did not noticeably impair EBV lytic cycle depletion of IgM and CD79A, suggesting that it is not required for BALF0/1 targeting of BCR in Burkitt cells, much of which resides at the plasma membrane (<u>Figure S5</u>E).

To further examine ubiquitin proteasome pathway roles in EBV lytic cycle BCR depletion, we tested whether BALF0/1 expression was sufficient to induce polyubiquitination of Ig heavy chain complexes. BALF0/1 or control BXLF1 cDNAs were expressed in human embryonic kidney (HEK) 293 cells, together with an EGFP-tagged IgM heavy chain construct. High molecular weight poly-ubiquitin conjugates were evident upon immunoblotting of immunopurified EGFP-IgM complexes from BALF0/1, but not BXLF1 expressing cells (Figure 4G). Furthermore, small molecule TAK-243 (49) inhibition of the E1 ubiquitin activating enzyme (UAE) ligase, which is the first enzyme in most host cell ubiquitin cascades, also significantly blocked BCR degradation in lytic-induced P3HR-1 cells (Figure 4H). These data suggest that BALF0/1 subverts the ERAD pathway to route internalized and polyubiquitinated BCR complexes to cytoplasmic proteasomes (Figure 4I).

### BALF0/1 BCR internalization and degradation requires the Ig heavy chain cytoplasmic tail

Since BALF0/1 is a cytoplasmic protein, we next characterized how it recognizes BCR complexes. Notably, CD79A and CD79B contain 61- and 48-amino acid cytoplasmic tails, respectively, whereas Ig heavy chains have as few as 3 cytoplasmic tail residues. We therefore hypothesized that BALF0/1 recognizes CD79A and/or CD79B cytoplasmic tails. To test this, we expressed sgRNA against CD79A or CD79B in P3HR-1 cells and then measured IgM abundance upon lytic reactivation. Despite substantial CD79A or CD79B depletion, EBV lytic induction readily targeted IgM for degradation (Figure 5A-B). However, as BALF0/1 may redundantly use either CD79A or CD79B to target BCR, we next depleted both CD79A and CD79B in P3HR-1. Upon lytic reactivation, EBV targeted IgM for degradation (Figure 5C-D). Similar results were observed by confocal microscopy, where IgM signals were nearly completely abolished in CD79A/B double KO cells induced for lytic induction. Bortezomib treatment resulted in colocalization between IgM and calnexin in 4-HT treated control and CD79A/CD79B double KO cells (Figure 6E-I and <u>S6</u>A-B). Thus, CD79 signaling chains are dispensable for BALF0/1-mediated degradation of membrane-bound immunoglobulin, much of which resides at the plasma membrane in latently infected Burkitt cells.

**Figure 5.**
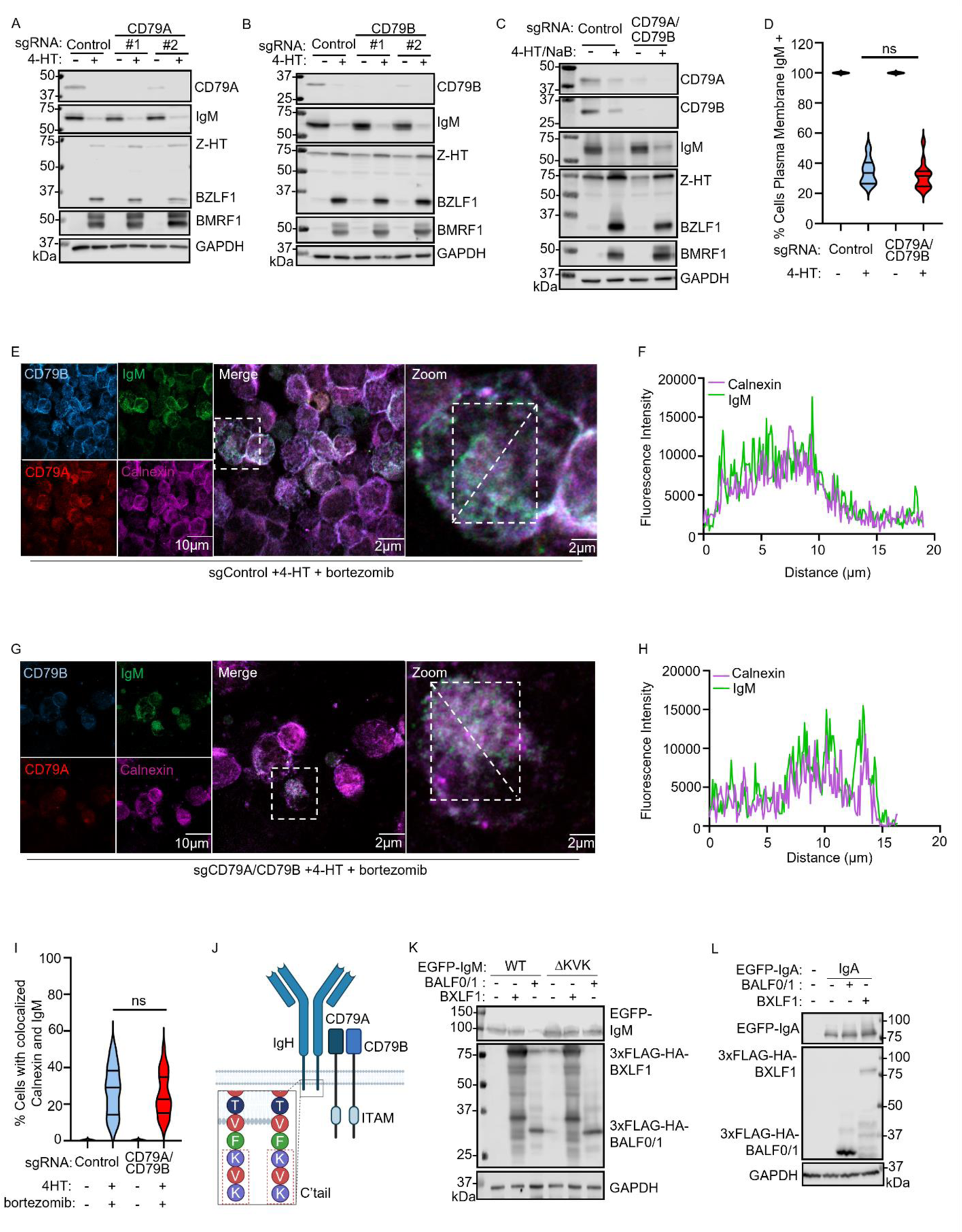
BALF0/1 mediated BCR degradation requires the Ig heavy chain cytoplasmic tail (A) Immunoblot analysis of WCL from Cas9+ P3HR-1 cells expressing the indicated CD79A targeting sgRNA, 4-HT induced into lytic cycle for 24 h. (B) Immunoblot analysis of WCL from Cas9+ P3HR-1 cells expressing the indicated CD79B sgRNA, 4-HT induced into lytic cycle for 24 h. (C) Immunoblot analysis of WCL from Cas9+ P3HR-1 cells expressing the indicated CD79A and CD79B sgRNA, 4-HT induced into lytic cycle for 24 h. (D) Mean ±SEM percentage of cells with PM IgM signals as in <u>Figure S6</u> of P3HR-1 cells expressing the indicated control or dual CD79A/CD79B sgRNA, uninduced or 4-HT induced, as indicated, using data from 12 randomly selected panels of 240 cells from n=3 replicates, analyzed using the ImageJ ComDet plugin. (E) Immunofluorescence analysis of CD79A, CD79B, IgM and calnexin in P3HR-1 cells expressing control sgRNA and 4-HT induced for lytic replication in the presence of bortezomib for 24h. (F) Fluorescence intensity line scanning for calnexin (magenta) and IgM (green) signals from the white rectangle region (E). (G) Immunofluorescence analysis of IgM, calnexin, CD79A and CD79B in Cas9+ P3HR-1 cells expressing CD79A and CD79B sgRNA. Cells were 4-HT induced for lytic replication and treated with bortezomib for 24 h. (H) Fluorescence intensity line scanning for calnexin (magenta) and IgM (green) signals from the white rectangle region in (G). (I) Mean ±SEM percentage of cells with overlapping calnexin and IgM signals as in panel E, G and <u>Figure S6</u> of P3HR-1 cells expressing the indicated control or dual CD79A/CD79B sgRNA, uninduced or 4-HT induced in the absence or presence of bortezomib as indicated, using data from 12 randomly selected panels of 240 cells from n=3 replicates, analyzed using the ImageJ ComDet plugin. (J) Schematic model of the BCR complex immunoglobulin heavy (IgH) chain, light chain and the CD79A and CD79B signaling chains. The C’ IgM cytoplasmic tail residues are shown. (K) Immunoblot analysis of WCL from 293T cells transiently expressing wildtype (WT) or cytoplasmic tail deleted (ΔKVK) EGFP-tagged IgM heavy chain alone or together with either BXLF1 or BALF0/1. (L) Immunoblot analysis of WCL from 293T transiently expressing the IgA heavy chain, alone or together with BALF0/1 or BXLF1. Statistical analysis was performed with Student’s t-test unless otherwise specified. ns p > 0.05. White bars indicate scale. See also <u>Figure S</u>6. Blots are representative of at least n=2 replicates.

**Figure 6.**
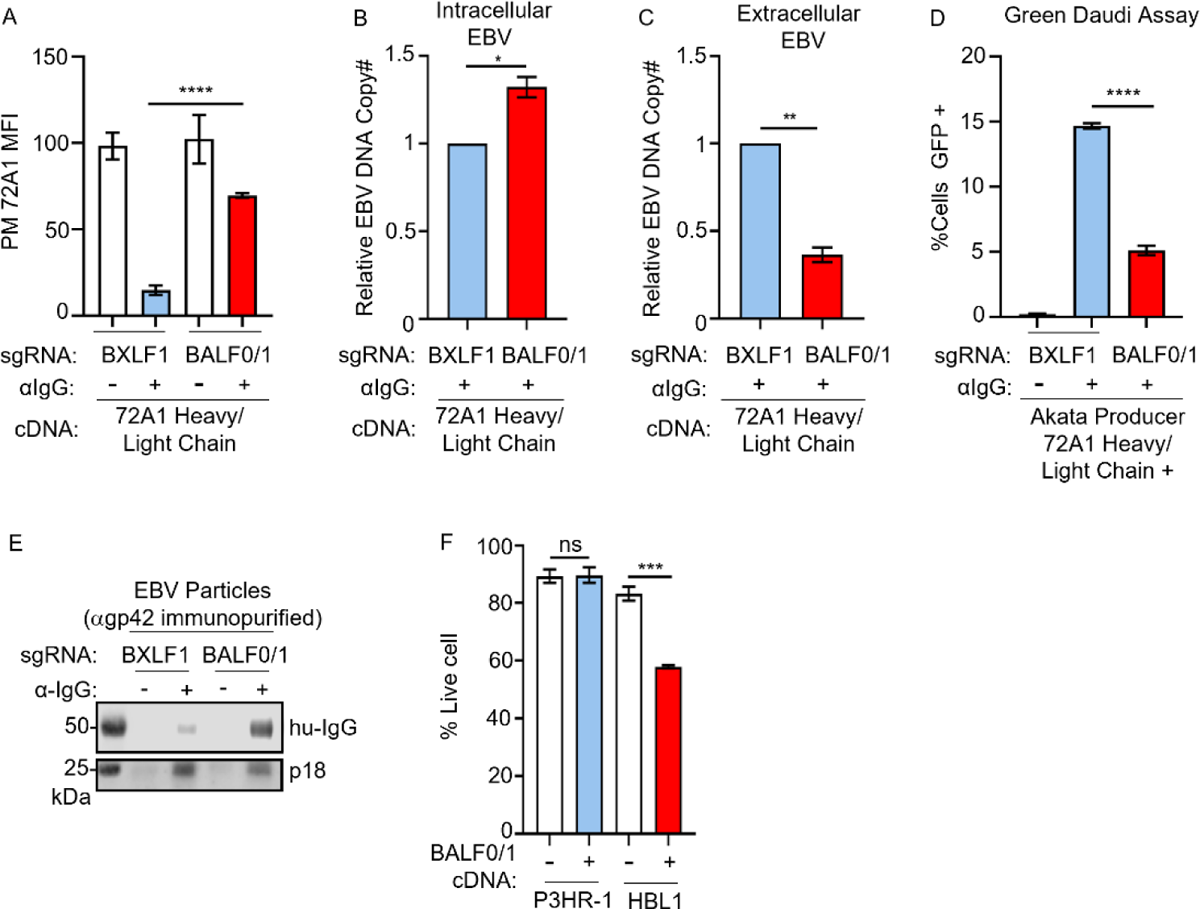
BALF0/1 supports virion release from cells expressing anti-EBV gp350 antibody (A) Mean ± SEM of PM 72A1 antibody abundance from n=3 replicates of Cas9+ Akata cells expressing the indicated sgRNA and stably expressing partially humanized, membrane bound anti-gp350 monoclonal antibody 72A1. The human-murine chimera 72A1 was detected by Alexa Fluor™ 488 tagged goat anti-mouse IgG secondary antibody. See also <u>Figure S7</u>A. (B) qRT-PCR analysis of EBV intracellular genome copy number from EBV+ Cas9+ Akata cells with stable membrane-bound 72A1 antibody expression together with BXLF1 or BALF0/1 sgRNA, induced into lytic replication by BCR crosslinking for 24 h. (C) qRT-PCR analysis of EBV extracellular genome copy number from EBV+ Cas9+ Akata cells with stable membrane-bound 72A1 antibody expression together with BXLF1 or BALF0/1 sgRNA, induced into lytic replication by BCR crosslinking for 24 h. (D) Mean ± SEM from n=3 replicates of green Daudi assay analysis of infectious EBV titers from EBV+ Cas9+ Akata cells with stable membrane-bound 72A1 expression together with BXLF1 or BALF0/1 sgRNA, induced into lytic cycle by α-IgG crosslinking for 24 h. (E) Immunoblot analysis of lysates of EBV virion immunopurified from supernatants of Cas9+ EBV+ Akata that stably expressed 72A1 together with BXLF1 or BALF0/1 sgRNAs, induced for lytic replication by α-IgG cross-linking. Shown are blots for human IgG (hu-IgG). (F) Mean ±SEM percentages from n=3 replicates of live P3HR1 or HBL1 cells mock induced or induced for BALF0/1 for 24 h, as determined by trypan blue exclusion assay. Statistical analysis was performed with Student’s t-test unless otherwise specified. ****p < 0.0001. **p < 0.01. *p < 0.05. See also <u>Figure S</u>7.

We therefore hypothesized that the Ig heavy chain cytoplasmic tail must be critical for BALF0/1-mediated BCR degradation. Notably, the IgM and IgG heavy chains contain only 3- and 28 residue cytoplasmic tails, respectively, including a conserved KVK motif (Figure 5J). To test whether the IgM cytoplasmic tail was necessary for BALF0/1 targeting, we transiently expressed either wildtype (WT) or C-terminal tail deletion mutant (ΔKVK) IgM in 293T cells, together with either BALF0/1 or BXLF1 cDNA. Immunoblot analysis showed a clear reduction of WT, but not ΔKVK IgM in cells co-expressing BALF0/1, but not in cells expressing control BXLF1 cDNA (Figure 5K). Interestingly, the IgA and IgE cytoplasmic tails do not contain a KVK motif, and IgA levels were not diminished by co-expression with BALF0/1 (Figure 5L).

### BALF0/1 supports virion release from cells expressing an anti-gp350 antibody

Tegumented EBV capsids acquire their lipid envelope by budding into the Golgi or Golgi-derived secretory pathway vesicles (50). Therefore, maturing EBV virion travel through the same secretory pathway compartments as immunoglobulin, and in the absence of degradation, membrane-bound immunoglobulin is incorporated into the virion envelope. We therefore hypothesized that BALF0/1 may have evolved to target the immunoglobulin, in order to cope with the situation where EBV infects a B-cell with BCR reactive to EBV glycoproteins, which could prevent virion release via tethering particles to the plasma membrane or could induce clumping.

To test this possibility, we stably expressed a membrane-bound IgG isoform of the murine monoclonal antibody 72A1, which reacts with EBV gp350, in EBV+ Akata cells (51). We used a humanized 72A1 heavy chain, in which the murine constant region was replaced by human the human Fc residues (51). We established BXLF1 vs BALF0/1 edited 72A1+ Akata cells and triggered them for EBV lytic reactivation by Ig cross-linking. We then measured relative EBV intracellular and extracellular genome copy number at 24 hours post-stimulation, an early timepoint prior to the onset of cell death, which could otherwise non-specifically release virion. As expected, FACS analysis demonstrated significantly lower levels of plasma membrane 72A1 in BXLF1 KO than in BALF0/1 KO cells, as detected by antibody reactive with murine light and heavy chain (Figure 6A). Interestingly, at 24 hours post Ig-crosslinking, we observed increased intracellular and decreased extracellular EBV genome copy number in BALF0/1 KO cells, relative to levels in BXLF1 KO controls (Figure 6B-C). Similarly, BALF0/1 KO resulted in a 3-fold decrease in infectious virions released from Akata cells relative to BXLF1 KO control cell levels, as measured by co-culture of the Ig-crosslinked Akata supernatants with Daudi target cells (the green Daudi assay (52)) (Figure 6D). It is worth noting that BALF0/1 KO did not affect late gene gp350 expression, suggesting an intact lytic cycle (<u>Figure S7</u>A) or cell viability (<u>Figure S7</u>B), indicating that differences in extracellular viral load were not based on differences in live cell number. These data suggest that BALF0/1 facilitates virion release from infected cells that express Ig reactive with a viral glycoprotein.

To further test the hypothesis that BALF0/1 prevents incorporation of Ig into virion, we induced Akata cells expressing control BXLF1 versus BALF0/1 targeting sgRNA into lytic replication by Ig-crosslinking for 24h. We then immunopurified virus particles from cell supernatants via pulldown with an antibody specific for the EBV glycoprotein gp42. Immunoblot analysis of purified EBV revealed incorporation of human IgG into virus particles produced in BALF0/1 edited Akata, but to a lesser extent in BXLF-1 edited Akata (Figure 6E). Thus, an important BALF0/1 role may be to downmodulate incorporation of Ig into virus particles, which if specific for an EBV glycoprotein could induce clumping of secreted virus particles.

### BALF0/1 impairs survival of non-Hodgkin lymphoma dependent on BCR signaling

Several subtypes of non-Hodgkin lymphoma, including activated B-cell-like (ABC) and germinal center B-cell (GCB) subtypes of diffuse large B-cell lymphoma (DLBCL), mantle cell lymphoma and Burkitt lymphoma subtypes depend on constitutive BCR signaling for survival (53–58). We therefore hypothesized that BALF0/1 expression would be sufficient to trigger cell death of BCR-dependent B-cells. To test this hypothesis, we inducibly expressed BALF0/1 in HBL-1 diffuse large B-cell lymphoma cells, which are dependent on BCR signaling for survival (59). As a control, we expressed BALF0/1 in P3HR-1, which we did not find to be dependent on BCR components in a CRISPR screen (60). While BCR abundance was similarly reduced by BALF0/1 expression in P3HR-1 and in HBL-1 (<u>Figure S7</u>C-F), BALF0/1 significantly reduced HBL-1 but not P3HR-1 live cell number (Figure 6F). These results suggest that BALF0/1 expression likely impairs BCR signaling when expressed in lytic B-cells and raise the possibility that cell permeable BALF0/1 could be developed as a novel therapeutic strategy to target B-cell lymphoma dependent on BCR signaling.

## DISCUSSION

As a B-cell tropic herpesvirus, EBV most cope with large quantities of immunoglobulin as it replicates within and egresses from host B-cells. Here, we identified that the EBV early lytic protein BALF0/1 targets the BCR complex for degradation. Rather than routing membrane bound BCR to lysosomes, BALF0/1 instead subverts the caveolin-dependent endocytosis and ERAD pathways to target BCR for ubiquitin-dependent proteasomal degradation (Figure 7). To our knowledge, BALF0/1 represents the first example of a viral protein that targets BCR complexes for degradation, and is the first viral protein identified to sequentially utilize the caveolar and ERAD pathways.

**Figure 7.**
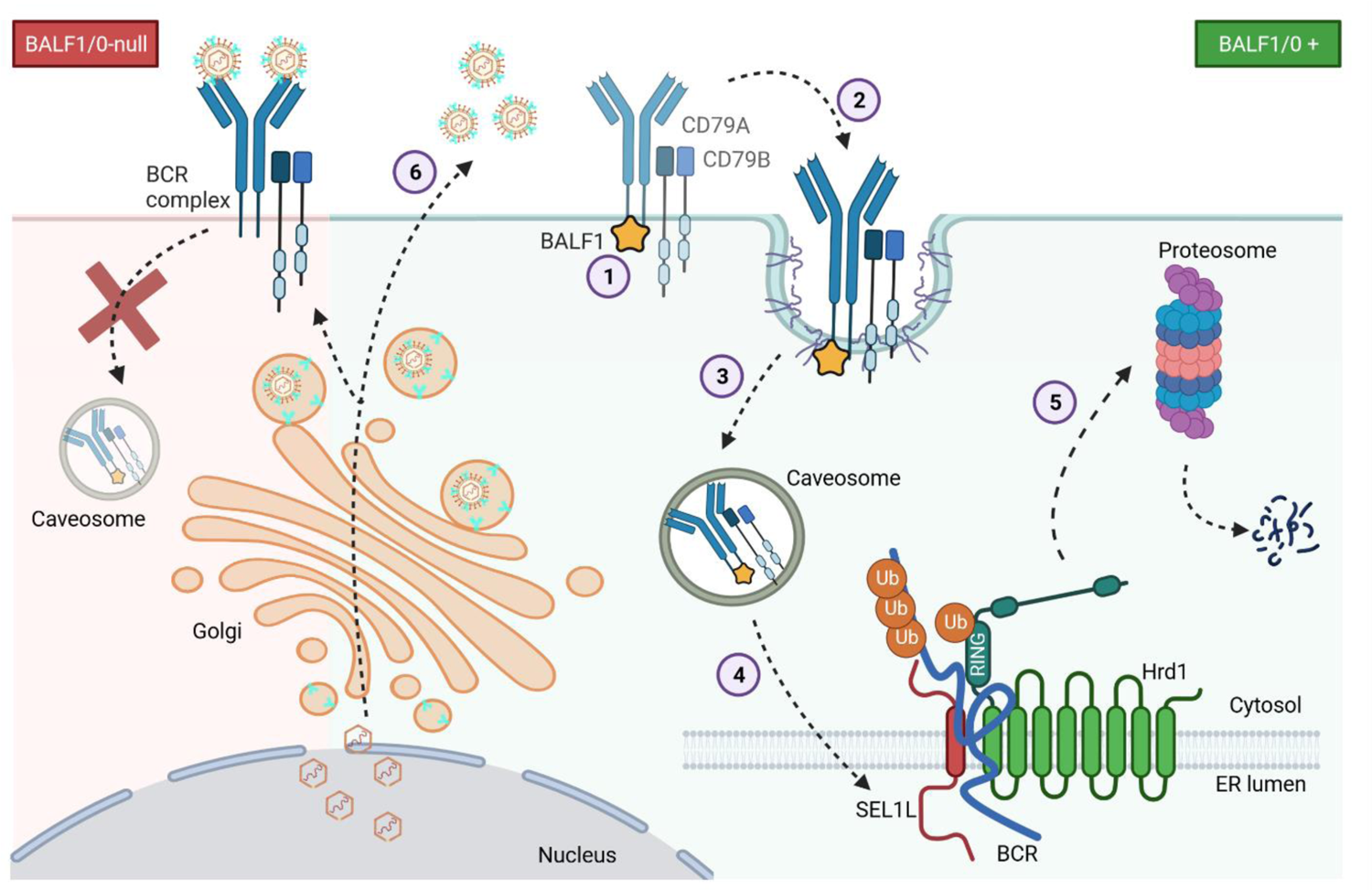
Schematic Model of BALF0/1 mediated BCR degradation Schematic model of BALF0/1-mediated BCR degradation. BALF0/1 recognizes the IgM heavy chain cytoplasmic tail and induces caveolin-1-dependent BCR endocytosis, delivering BCR to caveosomes. BCR are then retrograde trafficked to the ER, ubiquitinated and dislocated to the cytosol, where they are degraded by proteasomes. In the absence of BALF0/1, EBV release from B-cells with BCR reactive with virion glycoproteins is impeded.

We hypothesize that EBV evolved this strategy in order to facilitate release of virion from cells that express BCR reactive to an EBV virion component, which would otherwise impair virion assembly and/or release, for instance by tethering or causing clumping of egressing virus. In this manner, BALF0/1 may act analogously with human immunodeficiency virus (HIV) Nef and Vpu, which downmodulate CD4. This is necessary to support HIV release from CD4+ T-cells, since HIV utilizes CD4 as a major entry receptor, but must avoid being tethered by it as it egresses from CD4+ T cells (61–63). It would also be similar to viral evasion of the host restriction factor tetherin, which tethers viral particles to host cell plasma membrane to impair their release (64).

The observation that BALF1 targets BCR raises the question of how frequently EBV may infect B-cells reactive to viral glycoproteins. Germline encoded antibodies reactive to public viral epitopes, including multiple reactive with EBV gp350, were recently described (65). While EBV may encounter such B-cells at low frequency at the outset of primary infection, antibodies against virion structural proteins including gp350 and viral capsid antigen (VCA) are readily detected in EBV-infected individuals, suggesting the expansion of a pool of B-cells reactive with EBV antigens. In fact, anti-VCA antibodies are amongst the earliest detected upon primary EBV infection and are used to establish the diagnosis of mononucleosis. Anti-gp350 and VCA antibodies can also be abundant patients with EBV-associated cancers (66–71). As acute EBV infection progresses over the course of many months, it may be increasingly common for EBV to encounter a target B-cell with BCR reactive to viral components. For instance, we note that as many as 20% of tonsillar T-cells can be reactive with EBV encoded peptides in hosts with infectious mononucleosis (72), suggesting that high rates of EBV-reactive B-cells may be likely also be present. Furthermore, BCR reactive to EBV virion components may facilitate viral adhesion to and/or entry, potentially serving as novel co-receptors. Therefore, it may not be uncommon that EBV establishes latency in B-cells with BCR reactive to virion components.

B-cells invest considerable resources in immunoglobulin production. Another potential advantage of EBV lytic cycle degradation of high abundance BCR is that it may provide amino acid building blocks to support viral replication. It is less likely that EBV targets the BCR to prevent engulfment of viral antigens and their presentation to CD4+ T-cells, since EBV also reduces levels of MHC Class II in lytic cells (73). Notably, BALF0/1 is not known to be incorporated into the viral particle as a tegument protein (74). Therefore, in contrast to the major tegument protein BNRF1, which targets the SMC5/6 complex both in newly-infected and lytic cells (27), EBV instead evolved a mechanism to selectively deplete BCR upon lytic reactivation, but not in newly infected cells establishing latency.

It will be of interest to determine whether other γ-herpesviruses, including the Kaposi’s Sarcoma Associated Herpesvirus or murine herpesvirus 68, likewise evolved to target BCR for degradation. In support, multiple family members encode BALF0/1 homologs (75). However, since KSHV frequently superinfects EBV+ cells (76), it is plausible that KSHV relies upon EBV BALF0/1 to downregulate BCR, perhaps enabling the KSHV-encoded homolog to have evolved distinct functions.

BCR are internalized via clatharin-mediated endocytosis upon antigen binding (77–79), which delivers engulfed antigens to the lysosome for MHC Class II loading. EBV BALF1/0 instead triggers BCR internalization through a caveolar pathway reminiscent of the cholera toxin entry pathway (38). The principal structural component of caveolae, caveolin-1, is regulated by Src family kinase phosphorylation (80–85). Interestingly, our recent proteomic analysis identified that BALF0/1 associates with the serine/threonine protein phosphatase 2A (PP2A) regulatory subunit PP2R1A and also with the protein phosphatase 6 (PP6) catalytic subunit PPP6C (21). Since PP2A dephosphorylates Src kinases (86), and since the phosphatase inhibitor okadaic acid stimulates caveolae mobilization (87–90), we speculate that BALF0/1 may subvert host phosphatase as well as ubiquitin pathways to trigger BCR internalization.

While we did not obtain evidence that BAP31 is critical for BALF0/1 mediated BCR degradation, we speculate that it may nonetheless play an important role in particular contexts. For instance, it is plausible that another factor is redundant with BAP31 in Burkitt B-cells, but perhaps this redundancy does not exist in memory B-cells undergoing plasma cell differentiation, a key physiological site of EBV lytic reactivation for which in vitro models do not currently exist. BAP31 not only associates with BALF0/1, but it also co-purifies with B-cell receptors (43). Furthermore, multiple viral-encoded proteins that target host proteins for degradation also associate with Bap31, including Kaposi’s Sarcoma associated herpesvirus K3 and K5 (43, 91).

How does BALF0/1 trigger degradation of BCR complexes upon retrograde transport to the ER? Our data indicate that BALF0/1 stimulate ubiquitination of the IgM heavy chain cytoplasmic tail, and this is necessary for BCR degradation, since the E1 Ub ligase inhibitor TAK-243 rescued BCR abundance in the EBV lytic cycle. Since BALF0/1 is not known to have ubiquitin ligase activity and does not have homology to known ubiquitin ligases, our data suggest that it instead recruits an E3 ligase to slate immunoglobulins for ERAD. Thus, BALF0/1 may act as an adaptor or glue that juxtaposes BCR complexes with a host ubiquitin ligase to target BCR for degradation.

Why might BALF0/1 protein level targeting of BCR complexes be necessary in cells that also express the EBV alkaline nuclease BGLF5, which exerts host shutoff function beginning in the early lytic period (15)? First, immunoglobulin mRNA is highly abundant in B-cells, and a subset of BCR-encoding transcripts likely evades host shut-off. Second, even if host shutoff removes a large amount of BCR encoding transcripts, there is likely to be a large amount of assembling BCR complexes in the ER and secretory pathway and certainly also at the plasma membrane. Thus, BALF0/1 and BGLF5 may act synergistically to deplete BCR from cells undergoing lytic replication.

Multiple human lymphomas are dependent on BCR signaling for survival, including MCD DLBCL, chronic lymphomatous leukemia and subtypes of mantle cell and Burkitt lymphoma (53–58). Consequently, the BCR pathway Bruton’s tyrosine kinase serves as a major B-cell lymphoma therapeutic target, including by the small molecule inhibitor ibrutinib (92). BCR pathway kinase protein inositol phosphate 3 kinase delta (PI3K delta) inhibitors have also exhibited potent anti-lymphoma activity, though are limited by side-effects. Since knockdown of multiple BCR components is toxic to lymphomas dependent on BCR signaling (53, 54), our results suggest that BALF1 could potentially be leveraged in novel therapeutic approaches. For instance, chimeras between cell penetrating peptides, such as HIV TAT or antennapedia (93), with a potentially miniaturized version of BALF1 could potentially be delivered to lymphomas. Similarly, it might be possible to deploy cell-permeable BALF1 as an alternative to anti-CD20 monoclonal antibodies such as rituximab to limit autoantibody production.

In summary, we used a recently constructed EBV lytic cycle protein-protein interaction map to identify that the EBV early lytic protein BALF0/1 targets the BCR complex for degradation. BALF1 subverts the caveolin-1 pathway to internalize BCR complexes, which are trafficked in a retrograde manner to the ER, where they undergo proteasomal degradation via the ERAD pathway in a p97/VCP and ubiquitin dependent manner. The IgM cytoplasmic tail KVK motif, present also in the IgG cytoplasmic tail, was necessary for BCR degradation. BCR degradation facilitates virion release from B-cells encoding BCR reactive to a viral glycoprotein.

## METHODS AND MATERIALS

### Cell Lines

HEK293T were cultured in DMEM supplemented with 10% FBS and 1% Pen/Strep. P3HR-1-ZHT/RHT-Cas9+, EBV+ Akata-Cas9+, Raji-Cas9+, Daudi-Cas9+ and HBL1-Cas9+ cells were cultured in RPMI-1640 supplemented with 10% v/v FBS and 1% Pen/Strep. Cas9+ cells were maintained in 5 μg/ml blasticidin. EBV+-Akata-Cas9+ and P3HR-1-Z/R-HT-Cas9+ cells were also maintained with 25 µg/ml G418 and 25 µg/ml G418 and 25 µg/ml hygromycin, respectively. All cells were incubated at 37°C with 5% CO_2_ and were routinely confirmed to be mycoplasma-negative. Cell lines were authenticated by STIR profiling. Where inducible cDNA expression was used, cDNA were cloned into the pLIX-402 vector, which uses a Tet-ON TRE promoter to drive cDNA expression with a C-terminal HA Tag. Stable cell lines were generated by lentiviral transduction and antibiotic selection with puromycin. Cell lines were then maintained with 0.5 μg/ml puromycin.

### Chemicals and compounds

Unless otherwise specified, P3HR-1 ZHT/RHT or Akata cells lytic induction was triggered by addition of 400 nM 4-hydroxy tamoxifen (4-HT) (Sigma #68392-35-8) for 24 h or by 15 µg/ml α-human IgG (Agilent #A042301-2) for 48 h, respectively. 5 µg/ml doxycycline were used to induce BALF0/1 and BARF1 cDNA expression for 24 hours. Proteasome inhibitor bortezomib (5 nM, Sigma #5043140001), neddylation activating enzyme inhibitor MLN4924 (10 µM, Sigma # 5054770001), ERAD inhibitor eeyarestatin I (5-40 µM, Sigma #E1286), clatharin-mediated endocytosis inhibitor chlorpromazine (10-20 µM, Sigma # C8138), caveolae-mediated endocytosis inhibitor genistein (20-200 µM, Sigma # G6649), autophagy inhibitor bafilomycin A1 (10 µM, Sigma # 19-148), lysosomal protease inhibitors leupeptin (6.7 µM, Sigma #E18), E64 (10 µM, Sigma # E3132) or the E1 ubiquitin activating enzyme inhibitor TAK-243 (0.5, 1 and 2 µM, Selleckchem #S8341) were used where indicated. All inhibitors were added simultaneously with the lytic or cDNA stimulus.

### cDNA Expression Vectors

Unless otherwise specified, all cDNA were cloned into expression vectors by Gateway recombination. Briefly, 150ng of the donor vector containing the EBV cDNA and the destination vector were co-incubated with 1X LR Clonase Enzyme Mix (Invitrogen #11789-020) overnight at room temperature. The reaction mixture was then transformed into 50μl of Stbl3 *E. coli*, spread on LB plates with ampicillin. Entry vectors containing BALF1/0 and BXLF1 cDNA were gifts from Eric Johannsen. Entry vectors with IGHM and IGHA cDNA were obtained from DNASU (IGHM # HsCD00514313; IGHA # HsCD00512453 respectively). IGHM and IGHA were then cloned into the pDEST-CMV-N-EGFP, respectively (a gift from Robin Ketteler (Addgene plasmid # 122842; http://n2t.net/addgene:122842; RRID:Addgene_122842) (94). BALF1/0 and BXLF1 were cloned into PHAGE-3XFLAG-HA plasmid, respectively. To generate the IgM heavy chain cytoplasmic tail KVK deletion mutant (IGHMΔKVK), pENTR223-IGHM was used as a template for site-directed mutagenesis, using New England Biolab site directed mutagenesis kit (#E0554S) and the primers <u>Forward</u> 5’ GTCACCTTGTTCTAGGTGAAATGCCAACTTTC3’ and <u>Reverse</u> 5’ GAAAGTTGGCATTTCACCTAGAACAAGGTGAC 3’.

### Human-murine chimera 72A1 expression

cDNA for the anti-gp350 monoclonal antibody 72A1 heavy and light chains were gifts from Elliott Kieff and Fred Wang (51). To construct a partially humanized 72A1 vector, the murine 72A1 heavy chain variable region and light-chain variable region were previously cloned upstream of the human IgG1 constant region (together with a murine Ig tail sequence for membrane anchoring) and human Ig(κ) constant region of pEIG to create pCIRN-H1 and pEIG-L2, respectively. The murine-human chimeric heavy and light chain sequences were then cloned into pGenDONR by GenScript, with a P2A sequence added between the heavy and light chain sequences to create the pGenDONR-IgG H1-P2A-Ig(λ) L2 (tail) entry vector. Gateway LR cloning was used to shuttle this 72A1 cDNA into the TRC313 destination vector (Broad Institute) to create the TRC313-IgG H1-P2A-Ig(λ) L2 lentiviral expression vector. EBV+ Akata cells with stable 72A1 IgG H1-P2A-Ig(λ) L2 expression was generated by lentiviral transduction, hygromycin (50 µg/ml) selected for 2 weeks and then maintained with 25 µg/ml hygromycin. Detection of the murine-human chimera 72A1 detection was performed using goat anti-mouse IgG (H+L) cross-adsorbed secondary antibody Alexa Fluor™ 488 tagged (Invitrogen #A-11001).

### CRISPR analysis

CRISPR/Cas9 editing was performed as described (95). Briefly, sgRNAs were cloned into pLentiGuide-puro (a gift from Feng Zhang, Addgene plasmid #52963; http://n2t.net/addgene:52963; RRID:Addgene_52963)(96), pLenti-spBsmBI-sgRNA-Hygro (a gift from Rene Maehr (Addgene plasmid #62205; http://n2t.net/addgene:62205; RRID:Addgene_62205) (97) or LentiGuide-zeo (a gift from Rizwan Haq, Addgene plasmid #160091; http://n2t.net/addgene:160091; RRID:Addgene_160091)(98) and sequenced verified. Lentiviral transduction was performed as described previously (19). In brief, 293T cells were co-transfected with 500ng lentiviral sgRNA expression vector, 400ng psPAX2 (a gift from Didier Trono, Addgene plasmid #12260; http://n2t.net/addgene:12260; RRID:Addgene_12260) and 150ng VSV-G plasmids for packaging. Media was changed to RPMI at 24 hours post transfection 293T supernatants containing lentivirus were collected at 48 and 72 hours post-transfection, passed through a 0.45μm filter and transduced into P3HR-1-Z/R-HT-Cas9+, Akata-EBV-Cas9+ or HBL1-Cas9+ cells. Transduced cells were selected for 1 week with 0.5 μg/ml puromycin or 2 weeks with 25 μg/ml hygromycin or 100 μg/ml zeocin. CRISPR KOs were verified by western blot analysis. sgRNA against genes used in this study are as follows: ***BXLF1*** (5’ – TTGTAGTCCCTGAACCGATG – 3’); ***BALF0/1*** (5’ – CGGTCGAGGCGTGGGTCGC – 3’; 5’ – CATGTTCAGGGCCATGTACG – 3’); ***CAV-1*** (5’ – GTATTTCGTCACAGTGAAGG – 3’; 5’ – CGTTGAGGTGTTTAGGGTCG – 3’); ***Hrd1*** (5’ - TGAAGAGTGCAACAAAGCGG - 3’; 5’ – CTTGACTCACAAAGTCCACA – 3’); ***BCAP31*** (5’ – GAACCTCCAGAACAATCCCG – 3’; 5’ – CTTCATGTGGAAGTGCTCCA – 3’).

### Immunoblot analysis

Immunoblot analyses were performed as described previously (27). In brief, whole cell lysates were separated by SDS-PAGE, transferred onto nitrocellulose membranes, blocked with 5% non-fat dry milk in TBST buffer for 1 h and incubated with primary antibodies (1:1000) at 4°C overnight. Blots were washed 3 times in TBST and incubated with secondary antibodies for 1 h at room temperature. Blots were then washed 3 times in TBST solution, incubated with ECL chemiluminescence solution (Thermo Fisher #34578) and images were captured by Licor Fc platform. Antibodies used for immunoblot analysis in this study were: anti-Human IgM goat polyclonal antibody (Southern Biotech #2020-01), anti-Human IgG goat polyclonal antibody (Southern Biotech #2040-01), anti-HA-Tag (C29F4) rabbit mAb (Cell Signaling #3724), anti-GAPDH (D16H11) XP® rabbit mAb (Cell Signaling #5174), anti-CD79A rabbit polyclonal antibody (Proteintech #22349-1-AP), anti-CD79B (D7V2F) rabbit mAb (Cell Signaling #96024), anti-EBV BALF0/1 rabbit mAb (generated by Genscript for this study), anti-EBV ZEBRA Mouse mAb (BZ1) (Santa Cruz# sc-53904), anti-EBV Ea-D mouse mAb (1108–1) (Santa Cruz #sc-69679), anti-EBV p18 goat polyclonal antibody (Invitrogen #PA1-73003), anti-calnexin rabbit mAb (Cell Signaling #2433), anti-HRD1/SYVN1 rabbit polyclonal antibody (Proteintech #13473-1-AP), anti-BAP31 rabbit polyclonal antibody (Proteintech #11200-1-AP), anti-caveolin 1 mouse mAb (7C8) (Thermo Fisher #MA3-600), anti-GFP tag rabbit polyclonal antibody (Proteintech #50430-2-AP), goat anti-rabbit IgG, HRP-linked antibody (Cell Signaling #7074), goat anti-mouse IgG, HRP-linked antibody (Cell Signaling #7076) and bovine anti-goat IgG (H+L) HRP-linked antibody (Jackson ImmunoResearch Laboratory #805-035-180).

### Analysis of Ig heavy chain expression

For analysis of transient IgM and IgA heavy chain expression, expression vectors encoding for IgM heavy chain (IGHM), IgM heavy chain KVK deletion mutant (IGHMΔKVK) or IgA heavy chain (IGHA) cDNAs were transiently transfected into 293T cells, together with either BALF1/0 and BXLF1 cDNA, using Lipofectamine 3000, according to manufacturer’s instructions. Cells were harvested 24 h post-transfection and were washed twice with PBS. Cells was pelleted and lysed in cold lysis buffer (1% v/v NP40, 150mM Tris, 300mM NaCl in dH2O) supplemented with 1X cOmplete™ EDTA-free protease inhibitor cocktail (Roche # 11873580001), 1mM Na3VO4 and 1mM NaF for 1 h at 4°C with rotation. Lysates were precleared by pelleting insoluble debris and then incubated with 4X SDS loading buffer for 10 mins at 95°C.

### Analysis of human IgG in EBV virion

Supernatant containing EBV virion were collected from uninduced or lytic-induced (72 h) Akata-EBV-Cas9+ cells that express sgBXLF1 or sgBALF0/1. They were passed through a 0.45 µm filter and precleared with 2 μg/ml anti-rabbit isotype IgG antibody (Cell Signaling #2729) with protein A/G magnetic beads (Pierce, Thermo) for 1 h at 4°C with rotation. EBV virion immunoprecipitation was performed by incubating the precleared supernatant with 2 μg/ml anti-EBV BZLF2 (gp42) rabbit polyclonal antibody (Thermo Fisher # PA5-117635) for 4 hr at 4°C with rotation. Protein A/G magnetic beads were then added to the immunocomplex and were co-incubated at 4°C overnight with rotation. Beads were washed with lysis buffer four times and were eluted using 1X SDS loading buffer incubated for 10 mins at 95°C. Immunoblot analysis was performed and human IgG was detected by Goat Anti-Human IgM-UNLB (Southern Biotech #2020-01) antibody.

### Flow cytometry analysis

Cells were washed once with ice cold PBS supplemented with 2% volume by volume (v/v) fetal bovine serum (FBS). Cells were then incubated with the corresponding antibody at 1:500 in 2% FBS v/v, PBS for 30 mins at 4°C. Cells were pelleted, washed twice and resuspended in 2% FBS v/v PBS. Antibodies used for flow cytometry analysis performed in this study were: FITC-conjugated anti-human IgM (Southern Biotech #2020-02), goat anti-human IgG-UNLB (Southern Biotech #2040-01), Cy5-conjugated anti-EBV gp350 (clone 72A1) and donkey anti-goat IgG (H+L) Highly Cross-Adsorbed Secondary Antibody, Alexa Fluor™ Plus 647 (Invitrogen #A32849). For murine-human chimera 72A1 detection, goat anti-mouse IgG (H+L) cross-adsorbed secondary antibody Alexa Fluor™ 488 tagged (Invitrogen #A-11001) was used. For 7-AAD assays (Thermo Fisher, Cat#A1310), cells were harvested and washed twice with 1x PBS, supplemented with 2% FBS. Cells were incubated with a 1 μg/ml 7-AAD solution in 1x PBS / 2% FBS for five minutes at room temperature, protected from light. Cells were then analyzed via FACS on a BD FACSCalibur instrument and analyzed by FlowJo V10.

### Immunofluorescence analysis

Burkitt lymphoma cells dried on glass slides were fixed with 4% paraformaldehyde/PBS solution for 10 min, permeabilized with 0.5% Triton X-100/PBS for 5 min and blocked with 1% BSA/PBS for 1 h at room temperature. Cells were then incubated with a cocktail of primary antibodies (1:100 dilution) against CAV-1, IgM, IgG, calnexin, CD79A, CD79B, HA or EGFP in blocking solution for 1 h at 37°C. Antibodies used for immunofluorescence staining were the same as those for immunoblot analysis. Cells were then washed twice with PBS and incubated with a cocktail of secondary antibodies at 1:1000 in PBS for 1 h at 37°C in the dark. Finally, cells were washed twice with PBS and were stained with DAPI and mounted overnight with ProLong™ Gold Antifade. Image acquisition and analysis was performed with a Zeiss LSM 800 instrument and with Zeiss Zen Lite (Blue) software, respectively. Arivis Vision4D from ZEISS ZEN lite (blue edition) was used for 3D reconstruction in <u>Figure S2</u>C. Image J was used to score the % of colocalization of IgM-calnexin, IgG-calnexin, IgM-BALF0/1-HA, IgM-CAV-1 and IgG-CAV-1 using the ImageJ “Comdet” plugin.

### Immunoprecipitation and co-immunoprecipitation analysis

For poly-Ubiquitinylation analysis, 293T cells in 100mm cell culture dishes were co-transfected with 6000 ng each of plasmids expressing EGFP-IGHM and BALF1/0 or BXLF1 by Lipofectamine 3000 (Thermo #L3000001), according to manufacturer’s instructions. 5 nM bortezomib was added to the cells 6 h post-transfection. After 24 h, cells were trypsinized, harvested and washed twice with ice cold PBS. Pelleted cells were then lysed in ice cold lysis buffer (1% v/v NP40, 150 mM Tris, 300 mM NaCl in dH2O) supplemented with 1X cOmplete™ EDTA-free protease inhibitor cocktail (Sigma), 1 mM Na3VO4, 1 mM NaF, 1 mM PMSF, 4 mM 1, 10 o-phenanthroline, 2 mM sodium pyrophosphate and 1 mM EDTA for 1 h at 4°C with rotation. Lysed cells were pelleted and precleared with 2 μg/ml anti-rabbit isotype IgG antibody (Cell Signaling #2729) with protein A/G magnetic beads (Pierce, Thermo) for 1 h at 4°C with rotation. Precleared lysate was then incubated with 2 μg/ml anti-GFP antibody (Proteintech #50430-2-AP) for 1 hr at 4°C with rotation. Protein A/G magnetic beads were then added to the immunocomplex and were co-incubated at 4°C overnight with rotation. Beads were washed with lysis buffer four times and were eluted using 1X SDS loading buffer incubated for 10 mins at 95°C.

Where applicable, cDNA expression was induced by the addition of 5 μg/ml doxycycline. 100 million P3HR-1 cells were harvested and was lysed in ice cold lysis buffer (1% v/v NP40, 150mM Tris, 300mM NaCl in dH2O) supplemented with 1X cOmplete™ EDTA-free protease inhibitor cocktail (Roche # 11873580001), 1mM Na3VO4 and 1mM NaF for 1 h at 4°C with rotation. Lysates were pelleted and supernatants were incubated with anti-HA tag magnetic beads (Pierce, Thermo) at 4°C overnight. Beads were washed with lysis buffer four times and were eluted using 1X SDS loading buffer incubated for 10 mins at 95°C.

### EBV genome copy number quantification

For intracellular EBV DNA extraction, total DNA from 1×10^6^ cells was extracted using the Blood & Cell Culture DNA Mini Kit (Qiagen). Extracted DNA was diluted to 10 ng/μl and subjected to qPCR analysis, targeting the EBV BALF5 gene. For extracellular viral DNA extraction, supernatants from cells induced for lytic replication were collected and incubated with 25 mg/ml of DNase for 15 mins at 50°C, to degrade non-encapsidated DNA. Viral DNA was extracted by the Blood & Cell Culture DNA Mini Kit (Qiagen) and samples were subjected to qPCR analysis, targeting the BALF5 gene. For intra and extracelluar DNA analysis, serial dilutions of the pHAGE-BALF5 plasmid, beginning at 25 ng/μl, were used to generate a standard curve. Viral DNA copy number was then calculated by substituting sample Cq values into the regression equation dictated by the standard curve. qPCR primer sequences used for DNA copy number quantification are as follows: BALF5_forward 5’ GAG CGA TCT TGG CAA TCT CT 3’, BALF5_Reverse 5’ TGG TCA TGG ATC TGC TAA ACC 3’.

### Green Daudi assay

The green Daudi assay was performed as previously described (52, 99). In brief, EBV lytic replication was induced by anti-human IgG (15 µg/ml) crosslinking for 24 h in Akata producer cells that carry bacterial artificial chromosome EBV encoding a GFP transgene (100). Supernatant was collected and passed through a 0.45 µm filter. Daudi cells at 75,000 cells/ml were co-incubated with supernatant from EBV+ Akata producer cells. At 24 h post infection, culture media was exchanged to fresh RPMI, supplemented with 10% FBS. Cells were treated with 20 ng/ml tetradecanoyl phorbol acetate (TPA) and 3mM NaB for another 48 h. The percentage of GFP+ Daudi cells was then determined by flow cytometry.

### Quantification and statistical analysis

Unless otherwise indicated, all bar graphs and line graphs represent the arithmetic mean of three independent experiments (n = 3), with error bars denoting SEM. Significance between the control and experimental groups, or indicated pairs of groups, was assessed using the unpaired Student’s t-test in the GraphPad Prism 7 software. P values correlate with symbols as follows, unless otherwise indicated: ns = not significant, p > 0.05; *p % 0.05; **p % 0.01; ***p % 0.001; ****p % 0.0001. Pathway analysis was performed and visualized by using DAVID. Cytoscape version 3.8.1 was used to construct protein-protein interaction networks.

### Schematic Models

Biorender was used to create all schematic models.

## ACKNOWLEDGEMENTS

This work was supported by NIH RO1s AI164709, CA228700, U01CA275301 and R21 AI170751. We thank Dr. Domenico Tortorella for helpful discussions. M.P.W. is supported by the Medical Research Council (MR/W025647/1), Addenbrooke’s Charitable Trust, the Wellcome Trust Institutional Strategic Support Fund (204845/Z/16/Z) and the Cambridge Biomedical Research Centre, UK.

## Author Contributions

S.P.T.Y. performed and analyzed the experiments and bioinformatic analysis. S.P.T.Y., M.P.W. and B.E.G. supervised the study. S.P.T.Y. and B.E.G. wrote the manuscript.

